# Layer 6 corticothalamic neurons show diverse and dynamic responses that support a role in cortical gain control in noisy environments

**DOI:** 10.64898/2026.05.22.726967

**Authors:** Marina Cardoso de Oliveira, Patrick O. Kanold

## Abstract

Listening in complex environments requires the enhancement of relevant sounds and suppression of irrelevant ones. Corticothalamic neurons (CTNs) in auditory cortex layer (L) 6 provide modulatory feedback to thalamic nuclei and to upper cortical layers and are thought to be involved in gain control. However, their role in auditory processing remains unclear. We used *in vivo* two-photon imaging in mice to investigate the responses of L6 CTNs during active and passive listening. During a tone-in-noise detection task with varying levels of difficulty, L6 CTN responses exhibited higher gain in hit than miss trials, and this gain increased with task difficulty, indicating the active role of L6 CTNs. During passive listening, L6 CTNs exhibited diverse receptive fields and a heterogeneous tonotopic organization, with responses either suppressed or facilitated by background noise. Together, our results reveal diverse and dynamic context-dependent responses of L6 CTNs, consistent with a role in gain control.

## INTRODUCTION

One important feature of sensory processing is the enhancement of behaviorally relevant stimuli in the presence of irrelevant ones. For example, in the auditory system, the representation of attended sounds is amplified in the presence of background noise. Sensory cortices, including the auditory cortex, play an important role in this process, and computations within these circuits enable the robust encoding of stimuli in complex environments. However, it is still unclear how these circuits adapt to support such enhancements.

In the canonical cortical circuit, thalamocortical neurons primarily target layer (L) 4, which then project to L2/3, and subsequently to the deeper layers of the cortex, L5/6^1^. However, L4 is not the only thalamic-recipient cortical layer, as axons originating from the thalamus also send collaterals to L6^2–5^, while corticothalamic neurons (CTNs) in L6 provide feedback to thalamic nuclei^6^. Neural responses to sensory stimuli adapt to context and behavioral state^7,8^ to allow for encoding of sound stimuli across diverse environments, including in the presence of background noise. L6 CT circuits are hypothesized to play a role in this process^9^, as they integrate information from different areas and can modulate thalamic inputs to the cortex and upper cortical layers^10–14^. This modulation is thought to dynamically control the gain of sensory-evoked activity across all layers through disynaptic inhibition or monosynaptic excitatory connections in L4 and L5^15–17^, as shown by studies in the visual^12,13,18^, auditory^9^, and somatosensory cortices^14^. This dual pathway may support shaping the tuning curves of neurons in upper layers, contributing to robust sound perception in environments with high background noise.

Despite being well-positioned to integrate diverse sources of information, the role of L6, including L6 CTNs, in sensory processing remains poorly understood. For example, how L6 CTNs encode sound during auditory behavior is unknown. Optogenetic studies have shown that L6 CTNs can facilitate or suppress activity in upper cortical layers and the thalamus, depending on the timing of stimulation^14^; however, the functional significance of this modulation and how it is influenced by behavior remains unclear. Such dynamic modulation may be important for enhancing behaviorally relevant sounds while suppressing irrelevant ones, particularly in complex, noisy listening environments. Although auditory processing in the presence of noise has been extensively explored in the upper cortical layers^19–21^, studies of L6 have focused primarily on optogenetic manipulation of L6 CTNs^9,12,13,22^ and on identifying the sensory inputs that drive their activity^23^. Therefore, the functional properties of L6 CTNs in quiet and in noise, how these properties differ from those of upper cortical layers, and how L6 CTNs contribute to auditory perception across different behavioral states remain unknown.

We here set out to identify the functional properties and organization of L6 CTNs in the primary auditory cortex (A1) during passive and active listening and how the sound-evoked responses of L6 CTNs are modulated by background noise using *in vivo* two-photon imaging in mice. We found that under passive listening conditions, L6 CTNs comprise subpopulations with diverse excitatory receptive fields and prevalent inhibitory sidebands, showing disordered tonotopic organization. In the presence of background noise, L6 CTNs receptive fields changed, with responses being either facilitated or suppressed, depending on the subpopulation type. During a tone-in-noise detection task with varying signal-to-noise ratios (SNRs), L6 CTNs showed increased response gain on hit trials compared to miss trials, and this relative gain was inversely related to SNR. Together, our results show that L6 CTNs’ functional properties are consistent with a role of this cell population in cortical gain control. The heterogeneous tonotopic organization, together with the diverse FRAs and dynamic responses to background noise, suggests that these neurons are organized to differentially modulate the activity of upper cortical layers across different acoustic and behavioral contexts.

## RESULTS

### L6 CTNs exhibit sound-evoked responses and large-scale tonotopic organization

To selectively image L6 CTNs, we used the transgenic mouse line Ntsr1-Cre (B6.FVB(Cg)-Tg(Ntsr1-cre)GN220Gsat/Mmucd)^24^, crossed with a GCaMP6s expressing line^25^. To image L2/3 neurons, we used the transgenic mouse line Thy1-GCaMP6s (C57BL/6J-Tg(Thy1-GCaMP6s)GP4.3Dkim/J)^26^. We avoided the early onset of hearing loss associated with Cdh23^ahl/ahl^ in C57BL/6J mice by crossing founders with homozygous C57BL/6J-*Cdh23^Ahl/Ahl^* mice ^27^.

To characterize L6 CTNs’ responses to sound stimuli, we first performed widefield imaging^28^ to identify the anatomical location of A1 and the large-scale tonotopic organization of L6 CTNs (Figures 1A-1C). For that, we presented awake mice with five frequencies (4-64 kHz) at three sound-attenuation levels (50,70,90 dB SPL). Qualitative analysis of widefield fluorescence imaging showed that the response intensity to pure tones varies systematically along the tonotopic axis of A1, where low-frequency responses are stronger in the posterior-ventral region, and high-frequency responses are in the anterior-dorsal region of A1 (Figures 1B-1C).

**Figure 1:**
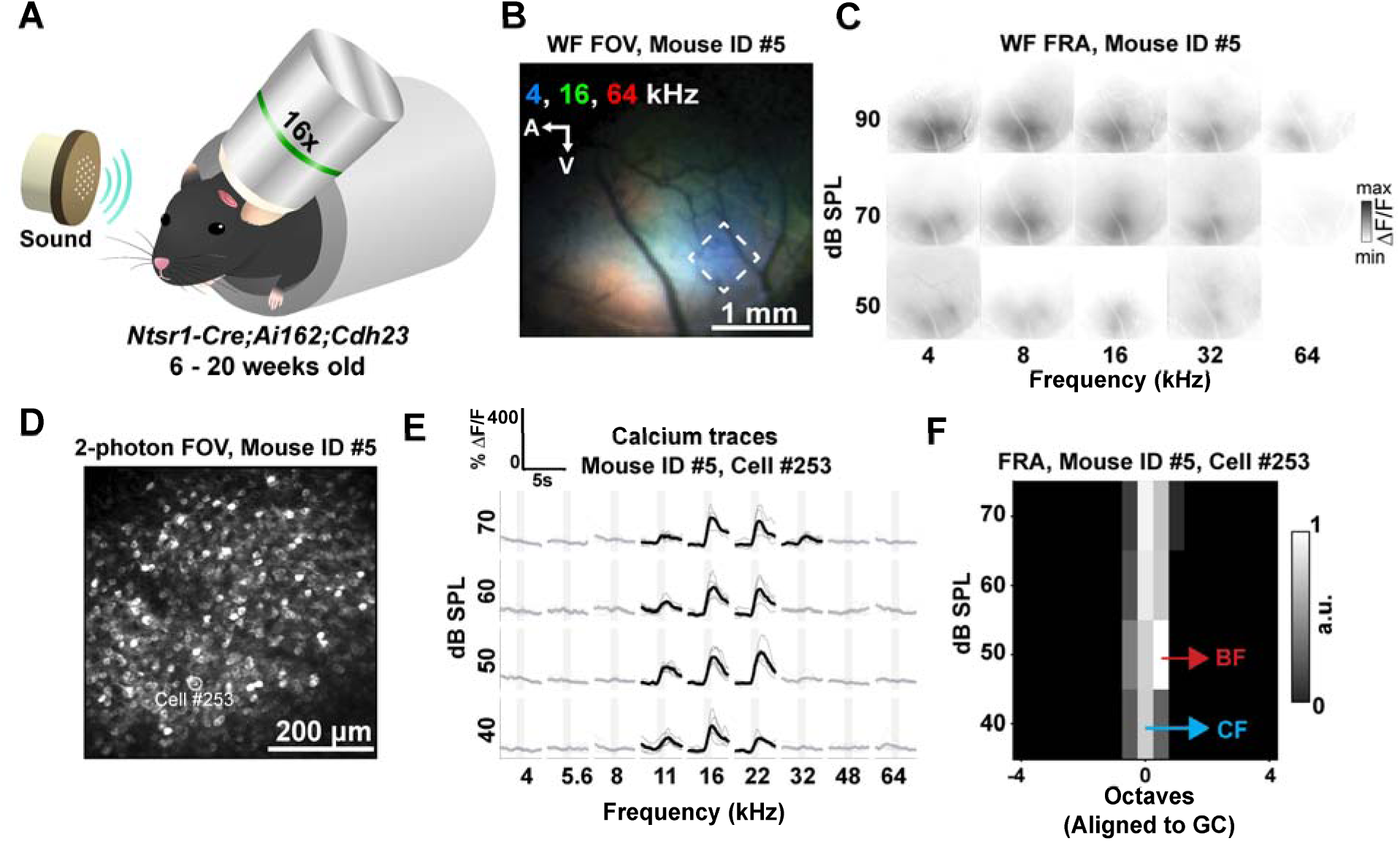
Functional *in vivo* imaging of layer 6 corticothalamic neurons in mouse auditory cortex. (A) Schematic of experimental setup for *in vivo* widefield and two-photon imaging. Imaging was performed in the left auditory cortex through a cranial window. Experiments were conducted in adult mice, 6 to 20 weeks of age, both males and females. 4x objective was used for widefield imaging and 16x for two-photon imaging. (B) Example of widefield fluorescence image of auditory cortex. Fluorescence responses to pure tones 4 (blue), 16 (green) and 64 kHz (red) highlight L6 CTNs large-scale tonotopic organization. Dashed area indicates the A1, where two-photon imaging was performed. (C) Plot of fluorescence changes over the widefield imaging field to varying frequency (x-axis) and sound levels (y-axis). (D) Example of field of view (FOV) (555 µm x 555 µm) of L6 CTNs at 770 µm deep in the cortex using two-photon imaging. Location corresponds to dashed area in (B). White circle highlights example neuron. (E) Example of sound-evoked fluorescence traces of sound-responsive CT neuron highlighted in (D) to tones (1s duration) of different frequencies and amplitudes. Bold black traces indicate frequency sound level combinations to which the cell was significantly responsive to the pure tone. Light gray traces indicate single trial responses. Bold gray traces indicate tones in which the cell did not show a significant sound-evoked response to the pure tone. Gray bar indicates stimuli presentation. (F) Frequency response area (FRA) of the example neuron in (E), aligned to the geometric center (GC) showing mean activity during 20 frames after three frames of each stimulus onset. BF indicates best frequency. CF indicates characteristic frequency. Responses are normalized to the maximum value.

### Tonotopy is not present at finer scales in L6 CTNs

We next performed *in vivo* two-photon imaging of A1 in awake passively listening Ntsr1-Cre mice (Figure 1D) (Z_Depth_ = 684 ± 62 µm, 17 mice, 22 FOVs) to investigate the sound-evoked responses of L6 CTNs to pure-tone stimuli. Mice were presented with nine frequencies (4-64 kHz) at four sound-attenuation levels (40-70 dB SPL) to characterize the FRAs of L6 CTNs. We identified 7787 L6 CT cells across 22 fields of view (FOVs) from 17 mice.

Previous studies have shown that neurons in L2/3 exhibit a heterogeneous organization of sound preference, while this heterogeneity is lower in L4^29–32^. To identify the tonotopic organization of L6 and how it compared to L2/3, we investigated the organization of sound preference in L6 CT cells by analyzing the distribution of frequencies on a spatial scale. For that, we calculated the best frequency (BF) and characteristic frequency (CF) of each significant sound-responsive neuron. The BF is defined as the frequency that produced the highest significant response regardless of the sound pressure level, while the CF is defined as the frequency that produced the highest significant response at the lowest sound pressure level (Figure 1F). The CF map of L6 CTNs (Figure 2A, left) appears heterogeneous and visually resembles a randomly organized CF map, in contrast to a more tonotopically organized L2/3 CF map (Figure 2B, left), which shows a stronger representation of low and high frequency areas.

**Figure 2:**
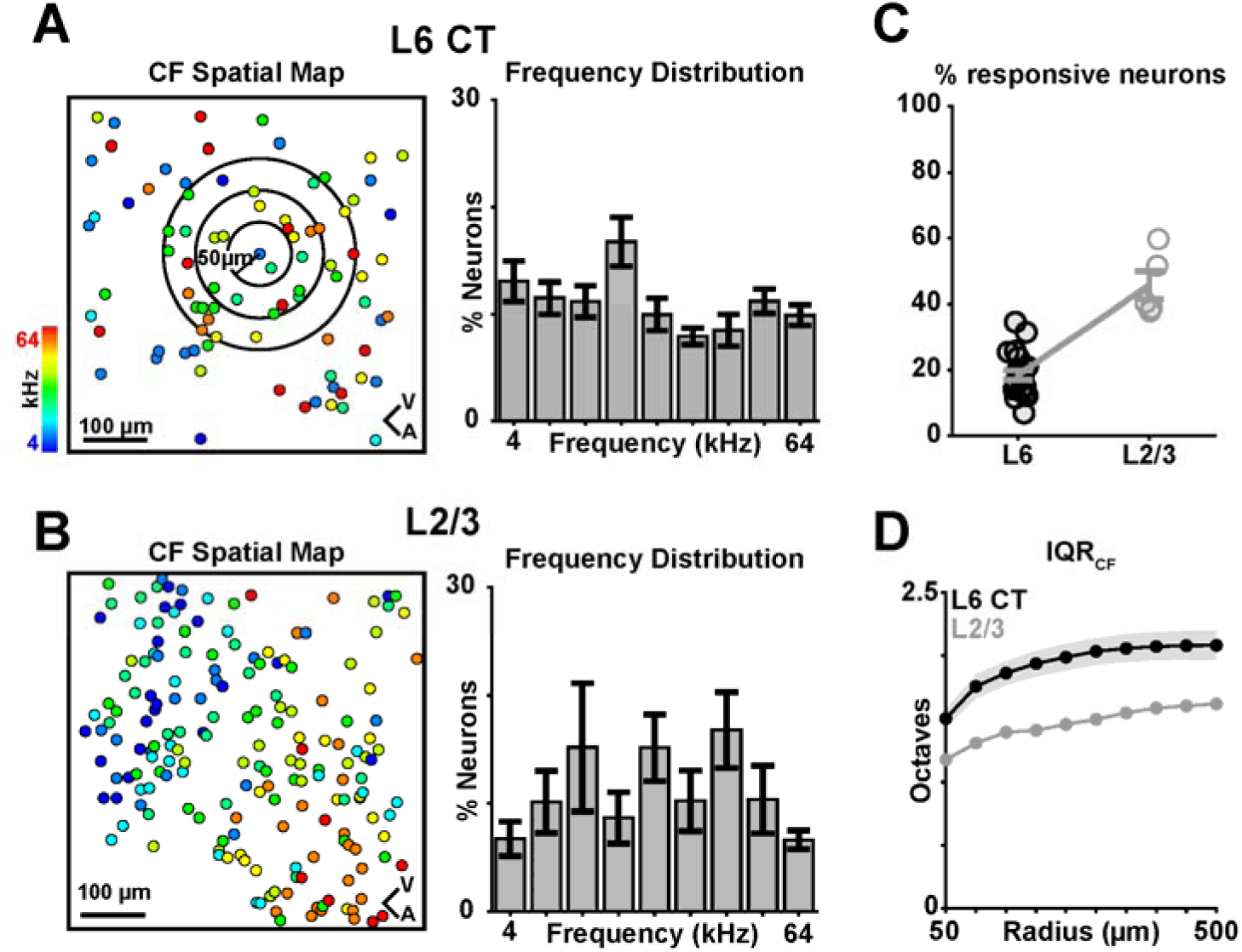
L6 CTNs show heterogeneous fine scale tonotopic organization. (A) Left: Example of spatial map of characteristic frequencies (CFs) for significant sound-evoked L6 CTNs in one FOV. Right: Proportion of CF values of significant sound-evoked L6 CTNs across different FOVs (n = 22 FOVs, 1329 sound-evoked neurons). Error bars indicate SEM. (B) Left: Example of spatial map of characteristic frequencies (CFs) for significant sound-evoked L2/3 neurons in one FOV. Right: Proportion of CF values of significant sound-evoked L2/3 neurons across different FOVs (n = 5 FOVs, 698 sound-evoked neurons). Error bars indicate SEM. (C) Percent of neurons in each FOV that are significantly responsive to at least one stimulus. Mean and standard deviation for each distribution: C57L2/3 = 0.29 ± 0.09, F1L2/3 = 0.28 ± 0.09, p > 0.68 (two-sample t-test); C57L4 = 0.30 ± 0.08, F1L4 = 0.31 ± 0.11, p > 0.91 (two-sample t-test). (D) CF variability (IQR_CF_) within different radius, for L6 CT and L2/3 neurons. Shaded area indicates SEM. Independent two-sample t-tests were performed at each distance bin after checking for normality (Kolmogorov-Smirnov), and it showed significant group differences from 50 to 300µm (p_50µm_ = 1.19 × 10^−2^, p_100µm_ = 9.94 × 10^−3^, p_150µm_ = 1.58 × 10^−2^, p_200µm_ = 1.09 × 10^−2^, p_250µm_ = 2.35 × 10^−2^, p_300µm_ = 3.31 × 10^−2^).

To quantify how frequency preference is organized in L6 CTNs, we analyzed the distributions of CFs within each FOV. We observed that CFs of L6 CTNs are mostly evenly distributed (Figure 2A, right). The peak we observe in the mid-frequency range of CF distributions (Figure 2A, right) is likely related to our choice of FOVs centered on A1, where mid-frequencies are overrepresented (Figure 1B). For comparison, we also examined the frequency distribution of CFs in L2/3 neurons, which showed a similar even distribution across the entire frequency spectrum, akin to that of L6 CTNs (Figure 2B, right).

We then quantified the fraction of sound-responsive L6 CTNs. 18.4 ± 6.7% of L6 CTNs show significant responses to at least one pure-tone stimulus, while 45.7 ± 9.5% of L2/3 neurons show significant sound-evoked responses to at least one stimulus (Figure 2C), consistent with previous findings in L2/3^30,33^. Thus, while L6 CT cells in A1 are sound responsive, fewer cells are responsive to pure tones compared with neurons in the upper layers of A1 under similar conditions (Figure 2C)^29,30^.

Following our observation on the heterogeneous spatial distribution of frequencies in L6, we evaluated the robustness of the tonotopic representation of sound frequency in L6 CTNs and compared it to L2/3. To analyze the spread of CFs for each FOV, we calculated the Interquartile Range (IQR) in octaves for all sound-evoked neurons that were within a range of 50 to 500 µm radius from each other^30^, as shown by the circles in Figure 2A, left. The IQR measures statistical dispersion and is defined as the difference between the 75^th^ and 25^th^ percentiles of the data. A higher IQR indicates greater heterogeneity. A tonotopically organized CF map would show lower IQR values for shorter distances and higher values for longer distances. The CF maps for L6 CTNs exhibit greater data spread and a more heterogeneous organization. In this case, neurons that prefer lower frequencies have a similar probability of being surrounded by neurons that prefer higher and lower frequencies (Figure 2D). In contrast, the IQR values for the CF organization of L2/3 neurons are lower and increase with distance, suggesting that neighboring neurons tend to have more similar frequency preferences, resulting in a lower spread of the data for shorter distances (Figure 2D). The IQR_CF_ of L6 CTNs is greater than that for both L2/3 and L4 under similar conditions^29,30^, and suggests that the CF organization of L6 CTN is more heterogeneous than L2/3 and L4. Our results suggest that tonotopic organization becomes more fragmented as auditory information travels the canonical cortical pathway.

### Receptive field shapes in L6 CTNs are diverse

Receptive fields in L2/3 of the mouse A1 can be classified into six distinct classes based on their spectral integration properties^33^. In L6 CT cells, tonal receptive fields have been unclear, as studies using different methodologies and species have shown contradictory results. Early *in vivo* patch-clamp studies in anesthetized rats suggested that L6 CTNs did not respond to tone^34^. Conversely, extracellular recordings in anesthetized cats found complex-shaped spectrotemporal receptive fields^35^, while two-photon imaging in awake mice found that L6 CTNs have well-defined V-shaped FRAs^9^. We therefore sought to investigate whether the FRAs of L6 CT cells fell into distinct classes beyond the V-shape. Since neurons in each FOV have different BFs, we aligned the FRAs to their geometric center as previously described^33^ and then applied a hierarchical clustering algorithm to the aligned FRAs. We selected the optimal number of clusters based on a visual inspection of the output dendrogram from the hierarchical clustering algorithm (Figures 3A, S1) and the corresponding clusters for each group. The optimal number of clusters was defined to maintain the maximum number that would represent different shapes, without repetition (Figure S1). Based on the dendrogram and the clustered FRA shapes, we found that the FRAs of L6 CTNs fell into six different shapes (Figure 3B), mirroring L2/3^33^. Classifying the FRAs into four or five different shapes results in visually distinct shapes being merged (Figure S1A-S1B), while using more than six shapes results in repetition (Figure S1C). Although the number of FRA shapes found is similar to those previously reported in L2/3^33^, they differ slightly in terms of type and distribution. L6 CTNs show a larger number of broadly tuned neurons (B-type and V-type) in comparison to sparsely tuned neurons (I-type and S-type) (Figure 3C). In contrast, L2/3 neurons are mostly sparsely tuned, as the majority of FRAs are I-type and S-type^33^. These analyses show that L6 CTNs form a diverse range of classes based on their FRAs, but that the classes differ from those seen in L2/3. These results also suggest that L2/3 and L6 CTNs differ in their associated circuits.

**Figure 3:**
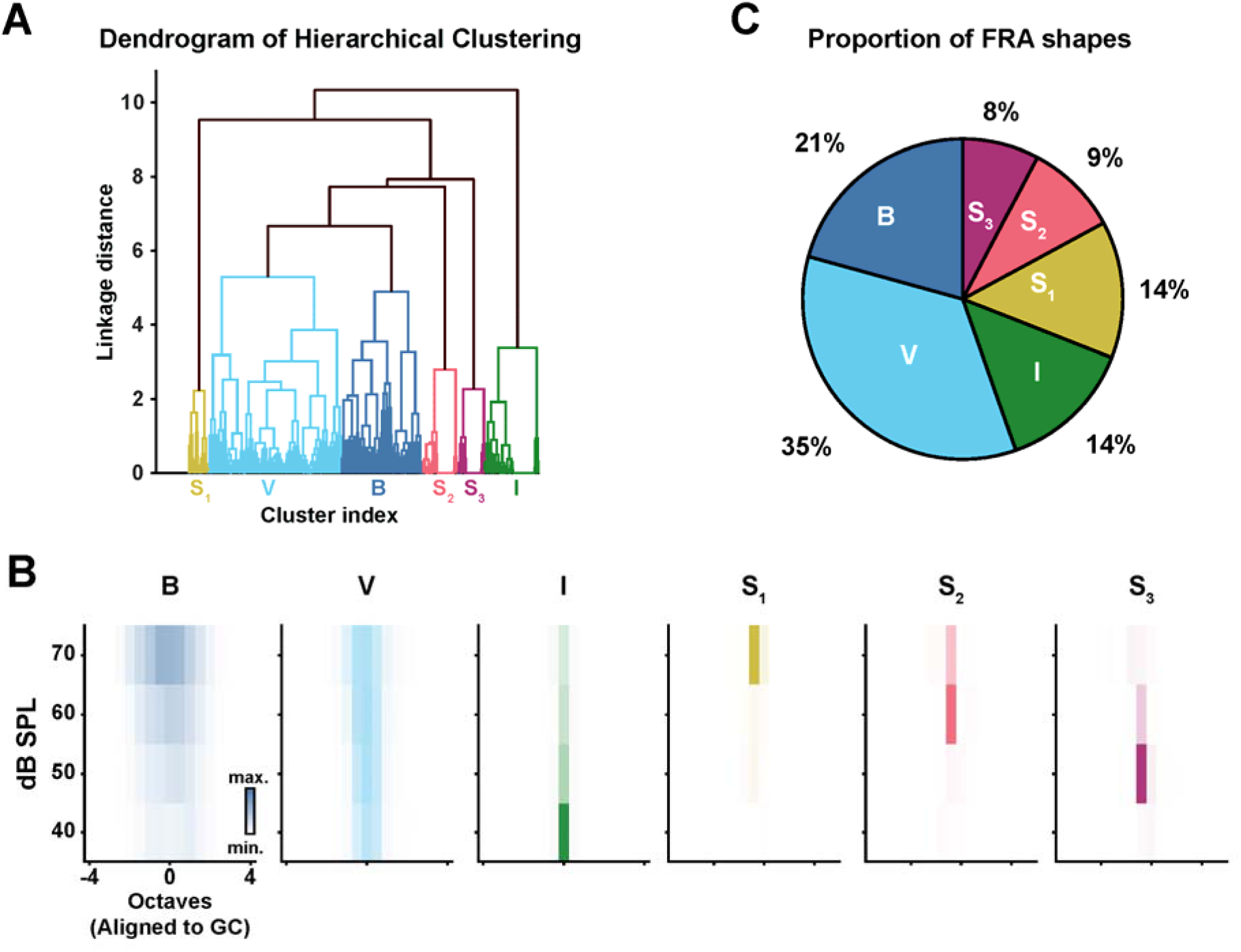
Unsupervised clustering of L6 CTNs responses reveals diverse frequency response area (FRA) shapes. A) Dendrogram of Hierarchical clustering, where each color represents an FRA shape. The linkage distance was one of the metrics used to define the optimal number of clusters (linkage distance = 5.5). B) Proportion of FRA types across 22 FOVs (n = 1142 neurons, 17 mice). C) Averaged FRA maps for each of the six clusters, aligned to the geometric center (GC). Clusters could be divided into Broadly tuned neurons, B-type; V shaped neurons, V-type; I shaped neurons, I-type; and sparsely tuned neurons, S1, S2 and S3. Number of clusters was defined by means of both visual inspection and linkage distance.

### Sideband inhibition of L6 CTNs is dependent on FRA shape

Inhibitory interactions play a crucial role in shaping neuronal responses and enhancing neuronal selectivity for specific stimuli. Neurons in the auditory system can exhibit inhibitory sidebands, enhancing the processing of stimuli with high spectral contrast^33,36–40^. To examine the inhibitory influences on the sound responses of L6 CTNs, we analyzed how the response to a neuron’s BF changes when a second tone is added to the stimulus.

We therefore presented both pure-tone and two-tone stimuli to passively listening awake mice. The two-tone stimuli were a linear sum of two pure tones at 60 dB SPL, resulting in a 63 dB SPL stimulus. We identified the BF of the neuron as the pure-tone frequency that elicited the strongest facilitated response and analyzed how the fluorescence traces (ΔF/F) changed when a second tone was introduced. We imaged 544 neurons that were significantly responsive to both pure-tone and two-tone stimulation (11 FOVs,11 mice). We found that L6 CTNs show both inhibitory and excitatory interactions (Figure 4A). When grouping the responses based on the clusters described above, we found that all clusters show wider inhibitory sidebands than excitatory tuning curves (Figures 4B-4C) (tuning width vs sideband width: p_B_ = 1.3 × 10^−3^; p_V_ = 1.1 × 10^−30^; p_I=_ 5.9 × 10^−32^, p_S1_ = 1.8 × 10^−22^; p_S2_ = 3.0 × 10^−35^; p_S3_ = 2.6 × 10^−10^, paired t-test with Bonferroni multiple comparisons correction). We also found that broader clusters (B and V-types) show narrower sideband width than the narrower clusters (I and S-types) (Figures 4B-4C) (sideband width: p_B,I_ = 8.7 × 10^−8^; p_B,S1_= 7.3 × 10^−7^; p_B,S2_= 2.3 × 10^−8^; p_B,S3_ = 2.1 × 10^−4^; p_V,I_ = 8.7 × 10^−14^; p_V,S1_= 1.9 × 10^−12^; p_V,S2_= 8.3 × 10^−15^; p_V,S3_ = 4.5 × 10^−8^; unpaired t-test with Bonferroni multiple comparisons correction), indicating relatively less sideband inhibition. Analysis of individual interactions within a single neuron revealed that the strength of interactions is similar for excitation and inhibition for most clusters, except for clusters V and I (p_V_ = 1.2 × 10^−3^, p_I_ = 1.5 × 10^−3^) (Figure 4D, left). However, there is net inhibitory than excitatory interactions (Figure 4D, right). To quantify the types of interactions in single neurons based on the FRA type, we calculated a Neuron Suppression Facilitation Index (SFI) (Figure 4E)^33^. An SFI of −1 or +1 indicates a purely facilitatory or inhibitory interaction, respectively. Consistent with our findings on individual interactions, B-type neurons exhibit the most facilitatory interactions, although these still represent a smaller proportion compared to inhibitory interactions. Despite the presence of broad inhibitory sidebands, particularly for I- and S-type neurons, the overall amplitude of inhibitory sidebands is smaller in L6 CTNs compared to L2/3 neurons^33^. Our results suggest that L6 CTNs show robust sideband inhibition, and that the broader tuning curves of B-type and V-type neurons are linked to reduced sideband inhibition when compared with I- and S-type neurons, consistent with a role for inhibition shaping the FRAs of L6 CTNs.

**Figure 4:**
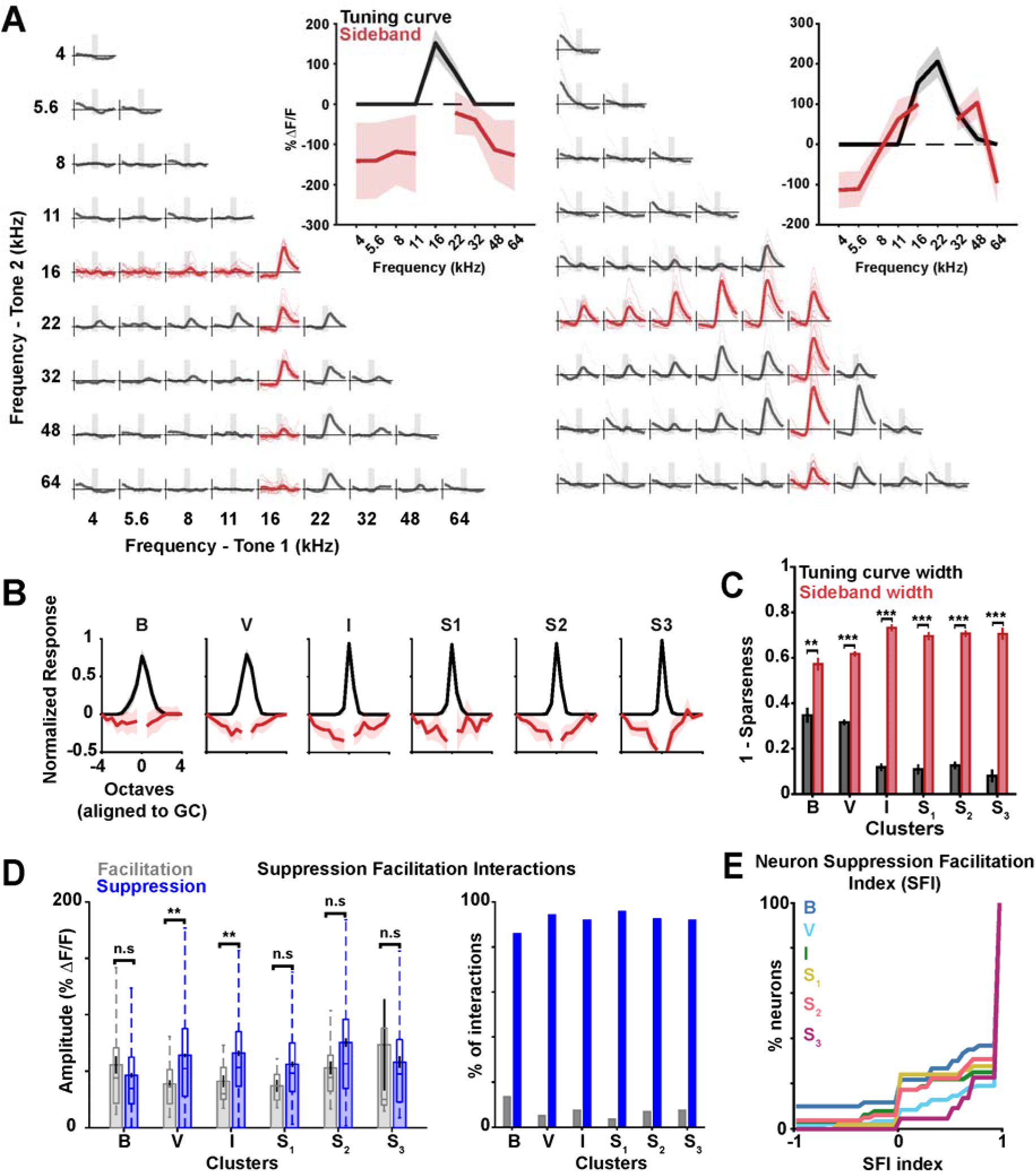
L6 CTNs show prevalent inhibitory interactions when a second tone is added to the neuron’s BF. (A) Example of fluorescence responses of two-tone stimuli in two representative L6 CTNs. Highlighted in red is the neuron’s BF and its respective responses to two tones. Inset shows the tuning curve (black trace) and inhibitory sideband (red trace) representative of the exemplar fluorescence traces. Shaded area corresponds to confidence interval. (B) Normalized mean tuning curves and sidebands for each cluster of cells, aligned to the geometric center (G.C.). Tuning curves were normalized to their maximum response. Sidebands were normalized to the maximum response of the tuning curve. Shaded area corresponds to SEM (n = 11 mice, 544 neurons). (C) Widths of the tuning curves (black) and the inhibitory sidebands (red) plotted as a function of FRA types. Inhibitory sidebands were significantly larger in width than those of the tuning curves, regardless of the FRA type (paired t-test with Bonferroni correction for multiple comparisons, tuning width vs sideband width ** p < 0.01,\*\*\**p* < 0.001). (D) Left: Amplitude of Suppression and Facilitation Interactions. Most FRA types show no difference between the amplitude of suppressed and facilitated responses, except for FRA types V and I. ** *p* < 0.01. Right: Proportion of facilitatory and suppressive interactions pooled over all neurons (n = 544 neurons) as a function of different FRA types. The percentage of suppressive interactions is considerably higher than facilitatory interactions regardless of the FRA type. (E) Suppression-Facilitation lndex (SFI) as a function of FRA types. Each line represents the cumulative sum of suppressive or facilitatory interactions within individual neurons. SFI = 1 corresponds to neurons that have purely inhibitory interactions, SFI = −1 corresponds to neurons that have purely facilitatory interactions.

### L6 CTNs show changes in functional properties when presented with broadband noise stimuli

The two-tone paradigm provides information about the inhibitory and excitatory components of the FRAs, but not how more complex stimuli, such as white noise, might affect the tuning and responsiveness of L6 CTNs. In low SNR conditions, the frequency tuning of neurons in superficial layers sharpens^21,41,42^. Furthermore, excitatory neurons, in supragranular layers more so than in the infragranular layers, exhibit greater suppression in response to broadband stimuli, such as white noise, than to narrower-bandwidth stimuli such as pure tones^43^ (Figure S1). This greater suppression can also be seen in the reduced strength of sideband inhibition in L6 CT compared to upper cortical layers, as observed in our study (Figure 4) and in previous reports in L2/3^33^. We thus sought to investigate how L6 CTNs’ responses to pure-tone stimuli would change in the presence of background noise with different SNRs. We imaged 3328 cells (11 FOVs, 10 mice), first with pure-tones only (PT), followed by pure-tones embedded in 50 dB SPL continuous white noise (PT+WN). Out of those cells, 950 neurons showed significant sound-evoked responses to PT and/or PT+WN. We found that more neurons are significantly responsive exclusively to PT+WN compared to PT (14.4% ± 5.5% responsive to PT and 26.8% ± 6.0% responsive to PT+WN), even though we observed a prevalence of inhibitory sidebands over excitatory sidebands. This could indicate that L6CTNs are more sensitive to more complex stimuli and non-linear interactions. We also observed that more than half of the population of responsive neurons is significantly responsive in both conditions (58.8% ± 8.3%) (p = 2.9 × 10^−2^, Wilcoxon Signed Rank test with Bonferroni correction for 3 comparisons). Thus, despite the presence of inhibitory sidebands, background noise unmasked tone-responsive neurons.

To investigate the influence of the background noise in more detail, we next compared the FRAs with and without noise. For the comparative analysis, we only considered neurons that are responsive to both PT+WN and PT (560 neurons). Neurons show diverse types of changes to their FRAs when background noise is added to the stimulus (Figure 5A). For example, broadly responding neurons with a wide FRA can become sparse in the presence of noise, indicating that the neuron now shows weak responses to fewer frequencies (Figure 5A, left). In another example, the FRA becomes sharper, and the neuron shows a narrower tuning and stronger responses to specific frequencies (Figure 5A, center). These types of changes would be consistent with the inhibitory sidebands we observed (Figure 4). However, in other cells in the presence of noise, the FRA became wider and showed stronger responses relative to the PT condition (Figure 5A, right) and effects consistent with the unmasking of tone-responsive neurons by noise. Thus, it seemed that background noise could have a suppressive or facilitative effect on the sound responses of L6 CTNs.

**Figure 5:**
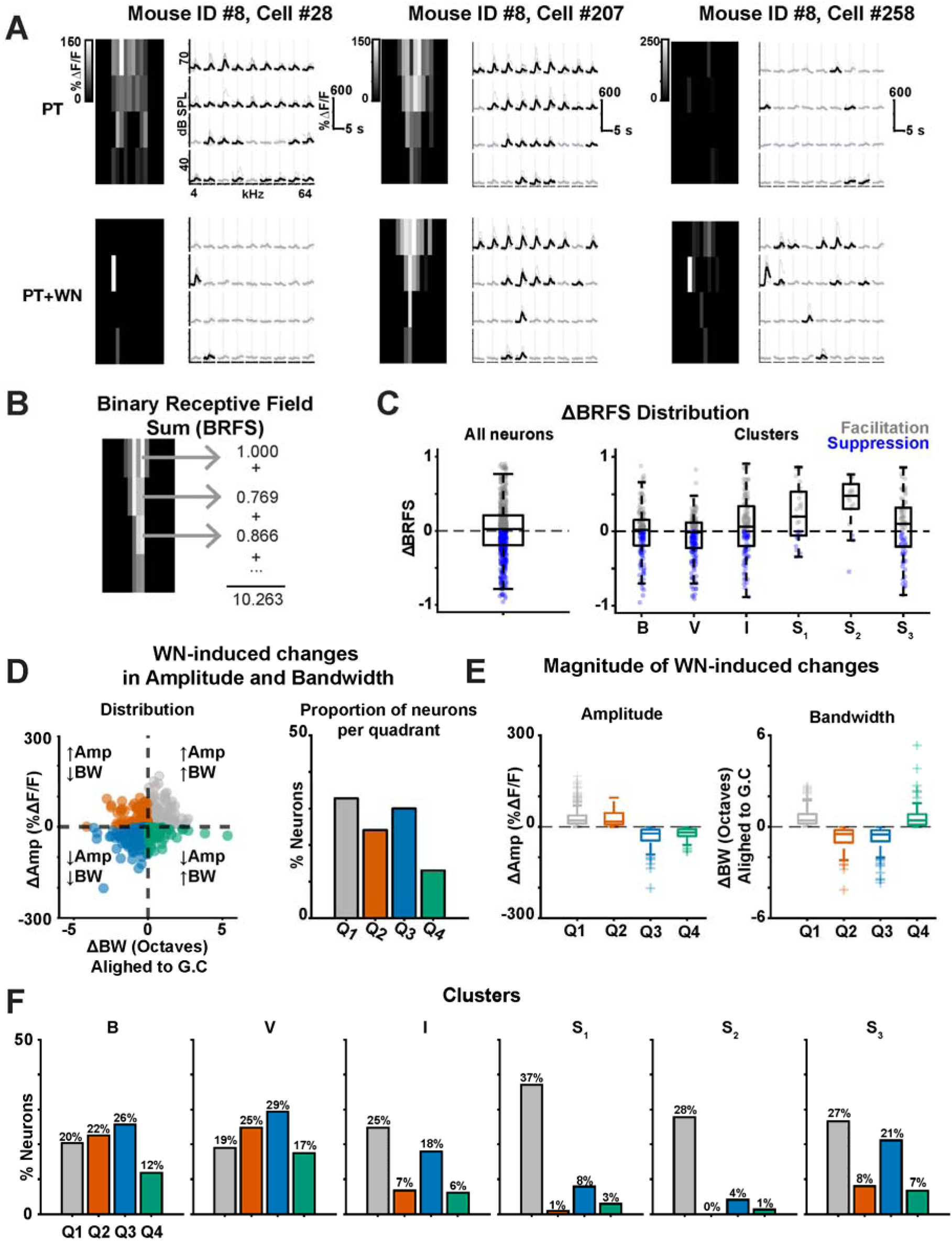
L6 CTNs FRAs show diverse types of effects when tones are played in the presence of background noise. (A) Examples of how FRAs of a single neuron can change in both PT (top) and PT+WN (bottom) conditions. Left: Example of FRA that becomes more selective once background noise is played. Center: Example FRA that becomes sharper and with stronger responses once background noise is played. Right: Example FRA that shows stronger and wider responses once background noise is played. (B) Example FRA illustrating the computation of the Binary Receptive Field Sum (BRFS) metric to assess how receptive field changes from PT to PT+WN condition. (C) Left: Distribution of ΔBRFS for all neurons that respond to both PTs and PTNs (n = 560 neurons). The proportion of neurons that show mostly suppressed responses (ΔBRFS < 0) is approximately 45.5% and the proportion of neurons that show mostly facilitated responses (ΔBRFS > 0) is approximately 54.5%. Right: Distribution of ΔRFS as a function of clustered FRA types. (D) Left: Scatter plot of differences in response amplitude (Amp) and bandwidth (BW) as a result of WN stimuli (11 FOVs, 10 mice, 560 sound evoked neurons). Right: Proportion of neurons in each quadrant of the scatter plot. (E) Magnitude of changes in Amp (left) and BW (right) as a result of WN stimuli. (F) Proportion of neurons that belong to different quadrants of the scatter plot separated by FRA-type.

To quantify these observed changes, we calculated the Binary Receptive Field Sum (BRFS) of each FRA (Figure 5B)^30^. This measure allows the comparison of the overall extent of responsiveness to sounds of various frequency/amplitude combinations. A lower BRFS indicates a selective response to a stimulus, while a higher BRFS indicates that a neuron is less selective and has a broader response area. To assess changes across different conditions (PT vs. PT+WN), we calculated the difference in BRFS between the two conditions (ΔBRFS = (BFRS_PT+WN_ - BFRS_PT_)/(BFRS_PT+WN_ + BFRS_PT_)). A ΔBRFS > 0 indicates that, for that neuron, in PT+WN there are mostly facilitatory interactions widening and strengthening the FRA, while ΔBRFS < 0 suggests that the interactions are mostly suppressive in PT+WN, yielding a smaller and weaker FRA. The ΔBRFS over the population, indicating an almost even proportion of neurons that are either facilitated or suppressed (mean ΔBRFS = 0.018 ± 0.36) (facilitated = 54.5%, suppressed = 45.5%) (Figure 5C, left).

Our two-tone analysis had revealed differences in sideband inhibition between neurons with broad and narrow FRA shapes. Thus, we next investigated if the direction of white noise-induced changes were related to specific FRA types. Given that cells with narrow (I and S type) FRAs have relatively broader sidebands than cells with broad FRAs (B- and V-type), we expected to see more suppression in narrow than in broad FRA types. However, the ΔBRFS distributions of different FRA types indicate that sparser FRAs (I- and S-types) are more likely to show facilitation in white noise than broader FRAs (B- and V-types), possibly because white noise could drive responses in regions of the receptive field that were not activated by pure-tone or two-tone stimuli, or due to nonlinear cross-frequency interactions that could be disinhibitory (Figure 5C, right).

To determine if changes in bandwidth or response amplitude drove those changes in ΔBRFS, we quantified how much each neuron’s amplitude and bandwidth changed under the PT+WN and PT conditions (Figures 5D-5E). Facilitated or suppressed responses could arise from changes in response amplitude, bandwidth, or both (Figure 5D). Most neurons exhibiting increased response amplitude in white noise also showed concurrent bandwidth broadening, indicating an increase in response gain and tuning width (Figure 5D). Also, reductions in response amplitude were typically accompanied by bandwidth narrowing. In contrast to findings from upper cortical layers^21,42^, white noise in L6 CTNs more frequently produced net facilitation than suppression (Figure 5D, right). Changes in the magnitude of response amplitude and bandwidth were greater for net suppressed responses than for facilitated responses (Figure 5E). When analyzing changes in bandwidth and amplitude across different FRA types, we observe that B- and V-type neurons more often exhibited suppression, while I- and S-type neurons tended to show more facilitation (Figure 5F), consistent with ΔBRFS analysis (Figure 5C). Together, these results indicate that L6CTNs exhibit diverse modulation when white noise is added to the stimuli. This modulation can manifest as different combinations of changes in response amplitude and tuning width, reflecting facilitation or suppression, depending on the FRA type. This increased facilitation in the presence of noise could be due to increased activity in L5 and disinhibition of inhibitory neurons in L6. In contrast, the suppression of L6 CTNs is likely due to the established inhibitory sidebands.

### L6 CTNs population activity is altered by background noise, whereas L2/3 neurons population activity is not

Previous studies have shown that single-cell responses in the auditory cortex are robust to a variety of stimulus parameters and background noise^7,44^ (Figure S2). Given that stimuli are represented by populations of neurons, we first investigated how the population activity of L2/3 is affected by background noise and if L6 CTNs showed similar changes. To assess population activity, we calculated pairwise neuronal signal-and-noise correlations between neurons in each condition (PT and PT+WN). We computed pairwise signal and noise correlations of all sound-evoked L2/3 neurons (5 FOVs, 4 mice, 211 ± 88 neurons/FOV) and found pairwise signal correlations did not differ between PT and PT+WN (Figure 6A), both when pooling all SNRs together (Figure 6A, left) or when separating by SNR (Figure 6A, right). Pairwise noise correlations also did not differ between PT and PT+WN (Figure 6B).

**Figure 6:**
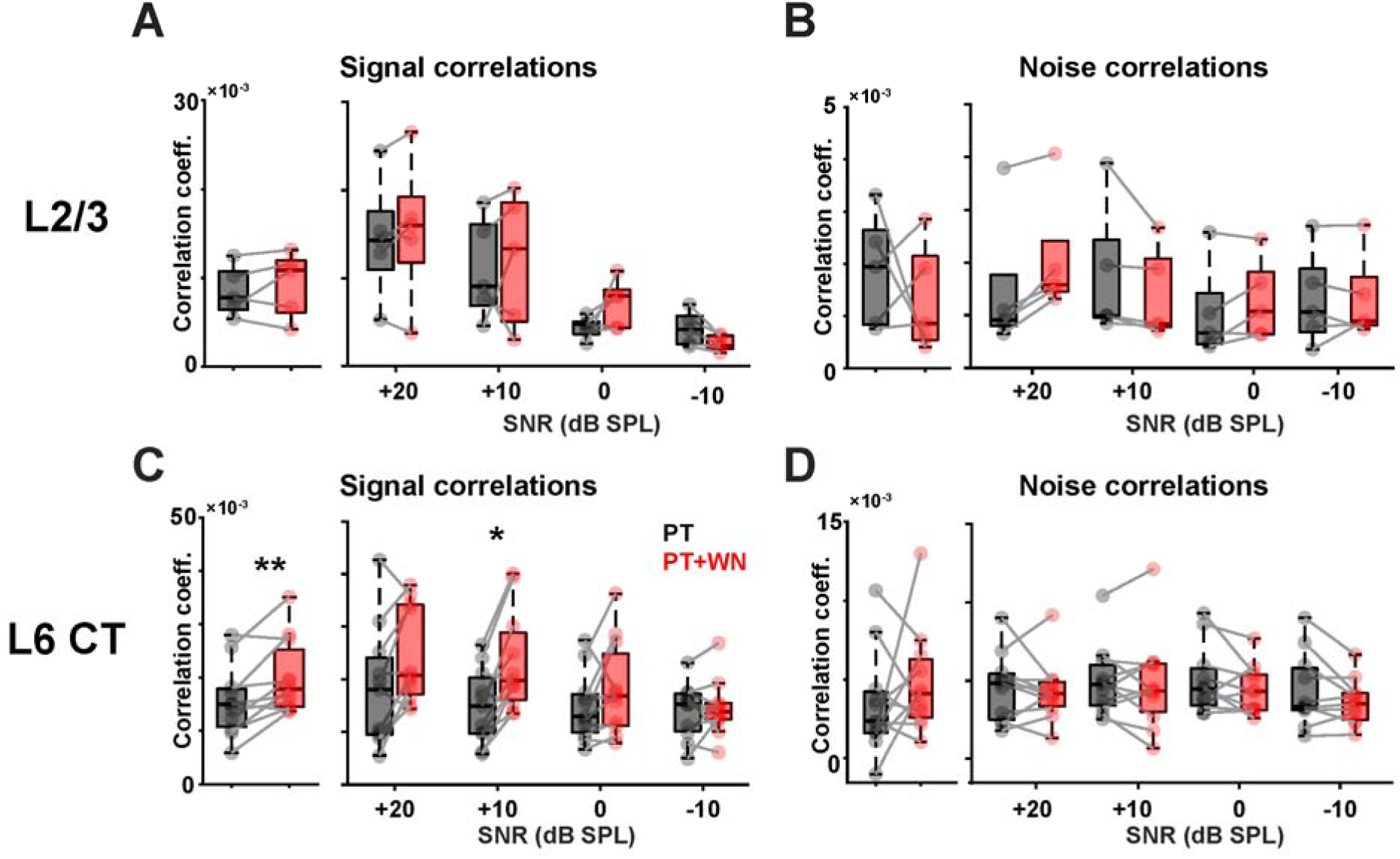
L6 CTNs but not L2/3 population activity differs between PT and PT+WN conditions. (A) Left: Plot of pair-wise noise correlations among L2/3 sound-responsive neurons in different conditions (PT in black and PT+WN in red) for all SNRs, (n = 5 FOVs, 4 mice). Two-sided Wilcoxon signed-rank test (p = 0.625) Right: Plot of pair-wise signal correlations among L2/3 sound-responsive neurons in different conditions as a function of SNRs. Two-sided Wilcoxon signed-rank tests with Bonferroni correction for multiple comparisons (*p_+20_* = 1, *p_+10_* = 1, *p_0_*= 0.75, *p_-10_* = 0.25) (B) Left: Plot of pair-wise noise correlations among L2/3 sound-responsive neurons in different conditions (PT in black and PT+WN in red) for all SNRs, n = 5 FOVs, 4 mice. Two-sided Wilcoxon signed-rank test (*p* = 0.812). Right: Plot of pair-wise noise correlations among L2/3 sound-responsive neurons in different conditions as a function of SNRs. Two-sided Wilcoxon signed-rank tests with Bonferroni correction for multiple comparisons (*p_+20_* = 0.25, *p_+10_* = 0.25, *p_0_* = 1, *p_-10_*= 1). (C) Left: Same as (A), but for L6 CTNs, n = 11 FOVs, 10 mice. Two-sided Wilcoxon signed-rank test (*p* = 0.0068359). Right: Same as (A), but for L6 CTNs (*p_+20_* = 0.054688, *p_+10_* = 0.019531, *p_0_* = 0.40625, *p_-10_* = 1). (D) Left: Same as (B), but for L6 CTNs n = 11 FOVs, 10 mice. Two-sided Wilcoxon signed-rank test (*p* = 0.51953). Right: Same as (B), but for L6 CTNs (*p_+20_* = 1, *p_+10_* = 1, *p_0_* = 0.69922, *p_-10_* = 1).

We next investigated the population activity of sound-evoked L6 CTNs (11 FOVs, 10 mice, 86 ± 18 neurons/FOV). L6 CTNs show increased pairwise signal correlations to tones played in a white noise background (p = 6.8 × 10^−3^) (Figure 6E, left) when considering all SNRs together and we observed a consistent upward trend in correlations for SNRs above 0 dB SPL. However, only the signal correlations at +10 dB SPL SNR show statistically significant differences between the PT and PT+WN conditions (p = 1.9 × 10^−2^) (Figure 6E, right). We did not observe differences in pairwise noise correlations between PT and PT+WN conditions, either when pooling all SNRs (Figure 6F, left) or when analyzing correlations separately for each SNR (Figure 6F, right). This suggests that background noise does not substantially alter shared variability across neurons in L6. These findings indicate that while L2/3 neurons’ single cell and population responses are mostly invariant to background white noise, L6 CTNs show changes in single cell and population activity with background white noise.

### Tone-in-noise detection at low SNRs reveals opposing modulation of L6 CTNs

Since L6 CTNs are sensitive to background white noise stimuli, we asked whether they could help control the gain of upper cortical layers during behavior, thereby possibly improving the perception of foreground tones. We speculated that if L6 CTNs played such a role, then the activity of L6 CTNs should be modulated by task performance and SNR levels. To investigate this possibility, we trained 4 mice on a tone-in-noise detection task in which they learned to turn a wheel in either direction when an 11.5 kHz tone was presented. The tone was played at different attenuation levels (30-70 dB SPL, 10 dB steps) over a continuous 50 dB SPL background white noise, resulting in five different SNRs (−20 to +20 dB SPL SNR) (Figure 7A). As expected, mice became expert at detecting the tone at high SNRs, with detection performance decreasing as the SNR decreased (Figure 7B). Trials were flagged as early when mice moved the wheel past 10° before the sound onset, and as hits when mice moved the wheel past 25° 200 ms after the tone onset (Figure 7C). Once mice reached a good performance, about 80% hit rate at the highest SNR, we performed two-photon imaging of L6 CTNs while mice performed the task (4 mice, 9 imaging/behavioral sessions/FOVs, 656 L6 CT sound-evoked neurons).

**Figure 7:**
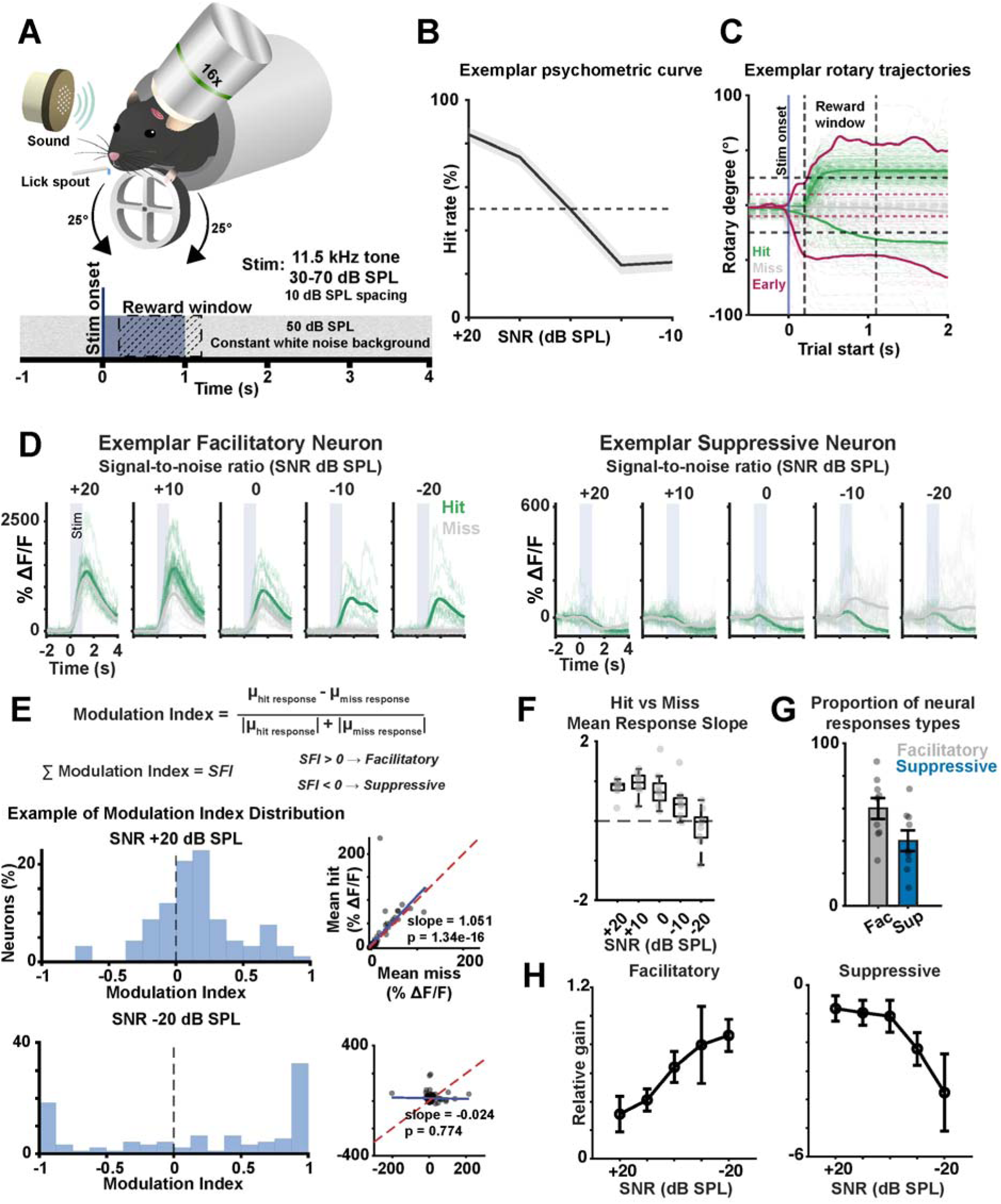
L6 CTNs show increased gain at low SNRs during a tone-in-noise detection task. (A) Behavioral paradigm of the tone-in-noise detection task. Mice were trained to detect a fixed 11.5 kHz tone presented at different attenuation levels (30-70 dB SPL) embedded in constant white noise (50 dB SPL). Mice reported tone detection by turning a wheel at the direction of their choice. Two-photon imaging was performed to record L6 CTNs during the behavioral task. (B) Example of psychometric curve for one animal averaged across 9 behavioral sessions. Shaded area represents SEM. (C) Example of rotary trajectories from one animal across 9 behavioral sessions. Hit thresholds are plotted as dashed horizontal gray lines, early thresholds are plotted as dashed horizontal red lines. The reward window is indicated by dashed vertical black lines. (D) Left: Example evoked responses from a neuron showing facilitation, with larger responses during hit trials (green traces) compared to miss trials (gray traces) at different SNRs. Right: Example evoked responses from a neuron showing suppression, with weaker responses during hit trials (green traces) compared to miss trials (gray traces) at different SNRs. The sum of modulation indices was used to calculate the suppression-facilitation index (SFI). (E) Top: Example distribution of the Modulation Index at the highest SNR, approximating a normal distribution (left). Linear regression of mean evoked responses at hit trials and miss trials (slope = 1.051, p = 1.34 × 10^−16^) (right). Bottom: Example distribution of the Modulation Index at the lowest SNR, approximating a bimodial distribution (left). Linear regression of mean evoked responses at hit trials and miss trials (slope = −0.0224, p = 0.774) (right). (F) Slopes of linear regression of mean evoked responses on hit and miss trials across SNRs (4 mice, 9 FOVs/sessions). (G) Proportion of facilitatory and suppressive neuros across behavioral sessions (4 mice, 9 FOVs/sessions), classified based on SFI. (H) Left: Relative gain of facilitatory responses, pooled based on SFI classification, at different SNRs (400 neurons across 4 mice, 9 FOVs/sessions). Right: Relative gain of suppressive responses pooled based on SFI classification, at different SNRs (256 neurons across 4 mice, 9 FOVs/sessions).

We first compared the sound-evoked activity of L6 CTNs depending on the behavioral outcome. L6 CTNs exhibit increased sound responses in hit trials compared with miss trials (Figure 7D), consistent with previous studies in upper cortical L2/3 neurons^8^. This increase in gain was observed in neurons that were facilitated by the tone (Figure 7D, left) and neurons that were suppressed by the tone (Figure 7D, right). The response difference between hit and miss trials seemed to be dependent on SNR, with high SNR showing only small differences. To quantify this effect, we calculated a modulation index of each neuron at each SNR (Difference between the median fluorescence response during hit trials and the median response during miss trials, divided by the absolute sum of the median responses) (Figure 7E). At high SNRs, the modulation index distribution was approximately normal, indicating that the calcium responses were largely similar between hits and miss trials (Figure 7E, top-right). Consistent with this observation, linear regression of the mean evoked responses during miss and hit trials resulted in a slope close to 1 (Figure 7E, top-left). In contrast, at the lowest SNR, the modulation index distribution approximates a bimodal distribution, indicating larger differences between miss and hit trial responses, and the slope of the linear regression was close to zero. The decrease in regression slope with decreasing SNR was consistent across behavioral sessions (Figure 7F), supporting the emergence of a bimodal pattern of the modulation index distribution at low SNRs, suggesting the presence of two distinct types of neurons in L6, one that becomes facilitated at hit trials, and another that becomes suppressed.

To classify neurons according to their response type, we then calculated the suppression-facilitation index (SFI), defined as the sum of the modulation index of each neuron across SNRs. Positive SFI values indicate neurons that were facilitated during hit trials, while negative SFI values indicate neurons that were suppressed during hit trials. Using this metric, we could separate L6 CTNs recorded during behavior into two groups (Figure 7G), with facilitatory neurons comprising most of the population. Separating L6 CTNs into these two groups showed response patterns that are dependent on SNRs (Figure 7H). In facilitatory neurons, the relative gain during hit trials increased as SNR decreased (Figure 7H, right). In suppressive neurons, this trend is reversed, with the relative gain becoming more negative as SNR decreased. Together, these results show that L6 CTNs are modulated by task performance, showing sound-evoked responses at low SNRs only when the animal detects the tone, consistent with a role for dynamic gain modulation in thalamus and upper cortical layers.

## DISCUSSION

In this study, we characterized how layer 6 corticothalamic neurons (L6 CTNs) in A1 are modulated by background noise and behavioral state. We find that L6 CTNs are sound-responsive, show a heterogeneous tonotopic organization, and have inhibitory sidebands. In the presence of a white noise background, many L6 CTNs showed increase in tone-responsive neurons and changes in their FRAs. During auditory tasks L6 CTNs dynamically regulate their activity in an SNR-dependent manner. These findings suggest that L6 CT cells are dynamically adjusting their responses to different listening conditions, which may enable them to modulate thalamic activity and influence upper cortical layers in response to changes in the auditory environment.

Neurons in the upper layers of the auditory cortex are modulated by stimulus context or attentional state^8,60–63^. Perception-dependent changes in L6 CTNs could play a role in this process. In a behavioral task, L6 CTNs responses exhibit higher cortical gain in hit trials compared to miss trials, and this gain increases with task difficulty. The presence of sound-evoked responses in hit trials at low SNRs indicates that the mouse can only perceive the sound when it is attending to the task, suggesting that L6 CTNs neurons are being modulated by the animal’s perception and state of attention^64–66^. This gain increase is consistent with L6 unique circuit topology, placing it in parallel to the canonical ascending pathway from thalamus →L4→L2/3 pathway, enabling L6 CT activity to directly modulate noise-robust responses in L2/3. Inhibitory circuits in A1 are important for separating background noise from behaviorally relevant stimuli^67^. Since L6 CTNs provide excitatory inputs to local inhibitory neurons that project to upper cortical layers^58^, L6 CT cells might contribute to the recruitment of inhibitory circuits that enhance cortical inhibition and improve the detection of stimuli in noise.

Genetically identified L6 CTNs are functionally diverse suggesting a variety of roles. We found that under passive conditions, L6 CTNs are responsive to sound and show a variety of FRA types. However, we found that the fraction of pure-tone responsive L6 CTNs is lower than in the upper cortical layers^29,30,33^. While previous studies on anesthetized gerbils^45^ and rats^34^ found that L6 CTNs are not robustly driven by sound stimuli, recent results in mice found that L6 CTNs exhibit well-defined FRAs^9,23,46^, consistent with our findings. The absence of sound-evoked responses reported in earlier studies may be due to anesthesia. L6 CTNs show a higher proportion of broadly tuned neurons (B- and V-type neurons) relative to sparsely tuned neurons (I- and S-type neurons) than L2/3 neurons^33^. The broader receptive fields found in L6 are consistent with findings in A1 of guinea pigs^47^ and cats^35^, as well as in the cat visual cortex, supporting the idea that deep cortical layers exhibit broader tuning across sensory modalities^3,48,49^. The broader frequency integration in L6 could arise from differences in thalamic or intracortical excitatory and inhibitory circuitry. In L2/3, strong inhibition from somatostatin (SOM) interneurons onto L2/3 pyramidal neurons contributes to sharp tuning and surround suppression^43,50,51^. In contrast, SOM interneurons may provide weaker inhibition onto L6 CTNs, potentially due to the higher threshold for depolarization of SOM interneurons in L6^52^, which could allow broader frequency integration in this layer. The two-tone stimuli paradigm^33,36,40,53,54^ revealed extensive inhibitory sidebands. This inhibition may arise from the activity of SOM interneurons that mediate the surround-suppression^43,55^.

L6 CTNs show a heterogeneous best-frequency map, coarser than that observed in L2/3 and L4^29,31^. This heterogenous tonotopic organization of L6 CTNs suggests that, although this layer receives inputs from collateral thalamocortical axons that target L4^3–6,56,57^, intracortical inputs from L6, L5, and L4 might play a key role in defining the response of these cells. This is consistent with weaker thalamocortical inputs to L6 CTNs than onto L6 inhibitory interneurons in the somatosensory cortex^58^.

Despite the presence of inhibitory interactions in L6, we find that, in contrast to L2/3, under passive conditions, background noise can have a net facilitatory effect on about half of L6 CTNs. In these neurons, background noise increases the response amplitude and broadens the FRA. In the other half, background noise suppresses response amplitude, and the neurons become more selective to specific frequencies. These changes suggest that L6 CTNs might differentially modulate the activity of upper-layer neurons in response to background noise. One group might become more specific to target stimuli (neurons that become more sharply tuned), possibly activating other excitatory neurons of similar tuning in L4 and L5^16,17^, while the other group (neurons that respond more broadly to tones in background noise) might engage inhibitory circuits^58^, including local cortical interneurons and thalamic interneurons in the thalamic reticular nucleus. Through this second pathway, L6 CTNs could contribute to suppression of neurons that are responsive to noise or non-relevant stimuli.

While population activity in L2/3 is more robust to background noise, pairwise signal correlations among L6 CTNs increase in background white noise relative to quiet conditions allowing these neurons to act synergistically. Taken together, our findings suggest that L2/3 shows mostly invariant responses to noise, while L6 responses show greater acoustic context-dependent changes consistent with a gain control role of L6 CTNs. This context-dependent variation of L6 CT activity might contribute to shaping noise invariance in the upper cortical layers of A1. Additionally, studies in the visual cortex have shown that L6 CTNs are involved in the gain control of upper cortical layers^12,13^, suggesting that these neurons may play an important role in extracting signals from background noise^59^.

In summary, we found that L6 CTNs are sensitive to the acoustic environment and are modulated in a perception-dependent manner, thereby potentially contributing to the emergence of a noise-invariant representation in the upper layers of A1.

## STAR METHODS

### Animals

All experiments and procedures were approved by the Johns Hopkins Institutional Care and Use Committee. For the passive experiments on L6 CTNs, we used 17 mice total (10 females, 7 males, 2 – 6 months). For the behavior experiment, we used 4 female mice (2 to 7 months between first training session and last imaging experiment). For L2/3 neurons experiments, we used 4 mice (3 females and 1 male, 2 to 6 months old). Mice were used based on availability. All mice were healthy and were only used for experiments detailed in this study. The following mouse lines were used: Ntsr1-cre (catalog #030648-UCD, Mutant Mouse Regional Resource Center; RRID:MMRRC_030648-UCD; ^24^), Ai162 (catalog #031562, The Jackson Laboratory; ^25^), *Cdh23* (catalog #002756, The Jackson Laboratory^68^), Thy1-GCaMP6s (C57BL/6J-Tg(Thy1-GCaMP6s)GP4.3Dkim/J)^26^. All mice used had C57BL/6J background. Ntsr1-cre mice were crossed with homozygous *Cdh23^Ahl/Ahl^*, to avoid the early onset of hearing loss associated with Cdh23^ahl/ahl^ in C57BL/6J mice^27^, and homozygous Ai162, which is a GCaMP6s reporter line, to generate heterozygous Ntsr1-Cre;*Cdh23*; Ai162 for experiments. Experimental mice were grouped housed with same sex littermates on a reversed 12hr light/dark cycle (dark from 9am to 9pm) and were provided food and water ad libitum. All experiments were conducted during the dark cycle.

### Cranial window and headpost implantation surgeries

Before the start of the surgery, mice were injected subcutaneously with dexamethasone (2 µg/g; VetOne) to prevent brain swelling during the cranial window implant. Mice were anesthetized with isuflorane (Fluriso, VetOne) using a calibrated vaporizer, induced at 4-5% and maintained at 1.5 - 2% for the duration of the surgery, and body temperature was maintained at 36.5°C with a feedback-controlled heating pad (Harvard Apparatus 50-7212). Eyes were covered with a thin layer of ophthalmic ointment to prevent drying (Bausch + Lomb Soothe). All surgical tools were sterilized prior to the first incision. Hair from the scalp was removed with a 5-minute application of Nair hair removal cream, followed by disinfection of the skin using 70% ethanol and betadine, applied consecutively four times. Surgical scissors were used to remove the skin and expose the top and left side of skull. Hydrogen peroxide was applied to dry and clean the skull. The skull was then scraped with scalpels to increase the surface area for better adherence of the headpost implant. Subsequently, muscles covering the left temporal bone were retracted to expose the left auditory cortex area. A craniotomy of a circular area of 4 mm diameter was performed above the left auditory cortex using a dental drill, and a stack of two round coverslips, one 3 mm layer (catalog #64–0720-CS-3R, Warner Instruments) stacked at the center of a 4 mm layer (catalog #64–0720-CS-3R, Warner Instruments) fixed with optic glue (catalog #NOA71, Norland Products), was placed on top of the exposed brain and secured with Superglue around the edge of the window. A customized headpost was fixed along the midline of the skull using Superglue, and dental cement (C&B Metabond) was added to improve headpost fixation and seal all exposed areas. After the surgery, mice were injected subcutaneously with Meloxicam (5mg/kg, MWI), and Cefazolin (300 mg/kg, West Ward Pharmaeceuticals) and kept on the controlled heating pad until major movements were observed. The same amount of Meloxicam and Cefazolin were injected at least 3 days after surgery. Mice were left to recover for two weeks before the start of the experiments.

### *In vivo* widefield imaging

Widefield imaging was performed to locate A1 within the cranial window as previously described^28^ Mice were head fixed on a customized station under an sCMOS PCO.edge camera with a 4x air objective (UplanSApo 4×/0.16, Olympus), mounted at an approximate perpendicular angle to the cranial window. The cranial window was then illuminated with a 470 nm LED (M470L3, Thorlabs). Images were acquired at 30 Hz, and both the camera and the sound stimuli were triggered externally (USB-6259, National Instruments). We played 5 pure tones from 4 to 64 kHz, at 3 different sound-attenuation levels (50-, 70- and 90-dB SPL) 10 times each to identify the sound responsive areas and different auditory subfields. The acquired images were analyzed as previously described^28^.

### Sound stimuli

Pure-tone stimuli (9 frequencies, 4-64 kHz, ∼ 0.5 octave spacing) were pre-generated with a custom MATLAB script that was loaded through a digital signal processor (RX6, Tucker-Davis Technologies) and programmable attenuator (PA5, Tucker-Davis Technologies). The generated waveforms were then played through an electrostatic speaker and driver (ES1, ED1, Tucker Davis Technologies) that was positioned approximately 10 cm from the animal’s right ear. The speaker was calibrated at 70 dB SPL reference with a Brüel and Kjær ultrasonic microphone and custom MATLAB scripts. Two-tone stimuli were pre-generated by attenuating each frequency to 60 dB SPL and then combining the two frequencies into a single waveform, resulting in a final attenuation level of 63 dB SPL. White noise stimuli were pre-generated by a custom MATLAB script and calibrated to 50 dB SPL reference with a Brüel and Kjær ultrasonic microphone.

### *In vivo* two-photon calcium imaging

Mice were head-fixed in a custom holder stage under an Ultima 2Pplus microscope (Bruker) paired with a tunable wavelength laser (Chameleon Discovery NX). For the experiments reported here, we used a 920 nm laser wavelength. A 16x objective (CFI75 LWD 16X W, Nikon) and galvo-resonance scanning were used to capture images at approximately 15 Hz and 1024×1024 pixel resolution at 1 to 2x magnification. *In vivo* two-photon calcium imaging was performed at the left A1, which was identified by the characteristic tonotopic axis observed through the pre-generated widefield imaging maps. The z-depth parameter was zeroed at the cortex surface, and the targeted imaging depth was approximately 680 µm (684 ± 62 µm) for L6 CT imaging and approximately 160 µm (161 ± 12 µm) for L2/3 imaging. Sound stimuli were triggered by the microscope’s frame-out signal.

### Data analysis

#### Two-photon calcium imaging data preprocessing

Image registration and fluorescence extraction from regions of interest (ROIs) were performed using Suite2p^69^. For L6 CT analysis, Cellpose was used to improve ROI detection^70^. For L6 CT analysis, all ROIs identified by suite2p were included and were further filtered using the sound-evoked criteria. For L2/3, only ROIs classified as cells were included in the analysis. The fluorescence signal used for analysis was calculated by subtracting the neuropil signal from the raw fluorescence signal (F= F_raw_ - 0.7 x F_neuropil_). We then calculated the change of fluorescence (ΔF/F) as (F - F₀)/F₀, where F_₀_ was defined as the median fluorescence value within a 225-frame window centered on the stimulus onset.

Response significance was determined based on prior methods^33^. Briefly, for each neuron, frequency, and amplitude combination, baseline fluorescence was defined as the mean ΔF/F within a 10-frame window immediately before stimulus onset. The response window was defined as the 20 frames following stimulus onset, with a 3-frame delay to ensure maximal separation of the response from baseline. For each window, the mean and 99.99% confidence interval (CI) were calculated using the Student’s t-distribution. A response was considered significant if the lower bound of the post-stimulus CI exceeded the upper bound of the baseline CI for at least one frequency–amplitude combination.

#### Interquartile range (IQR) analysis

To measure the variability within the field of view (FOV), we calculated the interquartile range (IQR) of the best and characteristic frequencies over different length scales (50 to 500 µm, 50 µm spacing) centered on each neuron. The IQR for each neuron was computed at each spatial scale and then averaged across all neurons to obtain the mean IQR per scale for each. This approach allows us to evaluate the distribution of frequencies within a specific area.

#### Hierarchical clustering classification of FRA types

We applied an unsupervised classification algorithm to identify distinct shapes of receptive fields within the L6 CT neuron population. The unsupervised classification was chosen to avoid human bias when selecting different FRA types. To avoid grouping cells based on their frequency preferences, rather than their shape, the FRAs were aligned to their geometric center, as described previously^33^. For each neuron, nonsignificant responses were set to zero, and the receptive field was normalized to the maximum response. A 20% threshold was then applied to remove the weakest responses and clean the FRAs for clustering. FRAs were then reshaped and zero-padded to a 1 × 68 matrix per neuron. We then used the MATLAB “linkage” function to cluster the normalized and thresholded FRAs. We used the Ward method and correlation distance to build the hierarchical clustering dendrogram. The optimal number of clusters was determined by visualizing the output dendrogram and the various clustering possibilities, selecting the number of groups that could best represent the maximum diversity of FRA shapes without repetition. Each neuron was classified according to different the FRA types. Finally, the pre-thresholding values were utilized to average the responses, resulting in the average FRA shapes for each cluster.

#### Two-tones experiment analysis

We identified the BF of the neuron as the pure tone that elicited the highest response and analyzed how the fluorescence traces (ΔF/F) changed when a second tone was introduced. The sideband frequency amplitude was calculated by subtracting the mean ΔF/F of the 20-frame post stimulus window (with a 3-frame delay) for the second tone from the mean value of the same window at the neuron’s best frequency. The tuning curve and sideband width were calculated as a measure of sparseness, similar as reported previously ^33^. Significant inhibitory interactions were identified when the mean ΔF/F of a sideband frequency was lower than the lower confidence interval (CI) of the tuning frequency response. Conversely, significant facilitative interactions were determined when the mean ΔF/F of a sideband frequency exceeded the upper CI of the tuning frequency response. Neurons that did not meet either criterion were considered to show no significant interaction. For each neuron, the Sideband Facilitation Index (SFI) was then calculated as: SFI = (S - F)/ (S + F), where S and F represent the mean ΔF/F values of the sideband and tuning frequency responses, respectively.

#### Tone and Noise experiments analysis

For the passive tone-in-noise experiments we first presented nine pure-tone stimuli (4–64 kHz) at four sound levels (40, 50, 60, and 70 dB SPL) under a no-noise (infinite SNR) condition (PT). We then added white background noise at 50 dB, resulting in SNRs of –10 to 20 dB SPL, at 10 dB SPL intervals (PTN). To quantify the changes in neuron’s FRAs at each condition, we calculated the binary receptive field sum (BRFS), as described previously^30^. Briefly, all non-significant responses in the FRA were set to zero and then the FRA was normalized to the maximum response before summing across all elements to obtain the BRFS. To assess changes in the receptive field shape for each experimental condition, we calculated the normalized difference in BRFS values (ΔBRFS = (BRFS_PT+WN_ - BRFS_PT_)/(BRFS_PT_ +BRFS_PT+WN_)).

To assess if the ΔBRFS changes were driven by white noise-induced changes in bandwidth (BW) or amplitude (Amp), we first calculated the geometric centered bandwidth of each neuron at each sound level. The bandwidth was defined as the tuning width at 60% of the best frequency, for each neuron and dB SPL. We then averaged the bandwidths across dB SPLs to obtain a single bandwidth value for each neuron under both conditions (PT and PT+WN). To calculate differences in bandwidth in PT+WN and PT (ΔBW), we subtracted the bandwidth for each neuron in the PT+WN condition from that in the PT condition. To assess differences in amplitude, we first identified the maximum response amplitude (%ΔF/F) for each neuron across different dB SPLs and experimental conditions. Next, we averaged these amplitudes to obtain a single maximum response value for each neuron and condition. To calculate ΔAmp we subtracted the maximum response amplitude in PT+WN condition from that in the PT condition.

#### Signal and noise correlations

Pairwise signal and noise correlations were computed based on previously reported studies ^27,29,30^. In summary, for the pairwise signal correlations, we averaged calcium responses across all repeats for sound-evoked neurons at each frequency and sound level. We then computed Pearson correlations between the mean calcium-evoked response over the defined response window (20 frames following stimulus onset, with a 3-frame delay) for every neuron pair, frequency, and sound level. Next, we averaged these correlations across frequencies and sound levels to obtain one correlation value per neuron pair and, finally, per animal. For the pairwise noise correlations, we averaged calcium responses across all repeats for sound-evoked neurons at each frequency and sound level. We subtracted the mean evoked trace from every trial. The resultant trace is then used to compute Pearson correlations between neuron pairs over a fixed time window of one second (15 frames) after the tone onset, for each frequency and sound level. Next, we averaged the pairwise noise correlations across frequencies, sound levels, and trials to obtain one correlation coefficient per neuron and, finally, per animal.

#### Behavioral experiments and data analysis

For the behavioral experiments, mice were weighed and water-restricted for 24 hours before the start of training. Our behavior paradigm was similar to that previously reported^71^. In summary, the speakers were calibrated to present an 11.5 kHz tone (1s) at different attenuation levels (30-70 dB SPL, 10 dB steps) over a continuous 50 dB SPL background of white noise, resulting in five different SNRs (−20 to +20 dB). The tone-in-noise detection paradigm consisted of the mouse spinning a rotor 25° between 0.2 s after the tone onset and 0.2 seconds after the tone cessation for a hit trial. If the spinning rotor was moved by at least 10° before the 0.2-second waiting period, the trial was marked as an early trial and not rewarded. If the mice waited for 1.2 seconds to spin the rotor after the tone onset, the trial was designated as a miss trial. Between each trial, 1 second with no rotor movement was required to allow the next trial to start. Mice were trained on this paradigm just enough to achieve about 80% hit rate at the highest SNR and then moved to perform the task during two-photon imaging. Each behavior session had at least 100 trials to be included in the analysis.

For the imaging data analysis during behavior, we defined sound-evoked responses following the same criteria as previously described for passive imaging analysis. Neurons were classified as sound-evoked if they presented significant sound responses at the highest SNR level.

### Statistical analysis

All statistical tests were performed in MATLAB R2025a. Detailed statistical information can be found in the figure legends. Data normality was using the Kolmogorov-Smirnov test. For data that did not follow a normal distribution, the Wilcoxon Signed Rank test was performed, followed by Bonferroni correction for multiple comparisons. For data that did follow a normal distribution, we performed paired t-tests with Bonferroni correction or N-way ANOVA with Tukey’s post-hoc test. For single comparisons, significance was established at p < 0.05. In cases of multiple comparisons, the Bonferroni correction was used to adjust p-values as necessary.

## Acknowledgements and Contributions

MCO and POK conceived the study. MCO performed experiments and analyzed data. MCO wrote the paper. MCO and POK edited paper. Supported by NIH RO1DC017785 (POK).

---

**Figure S1:**
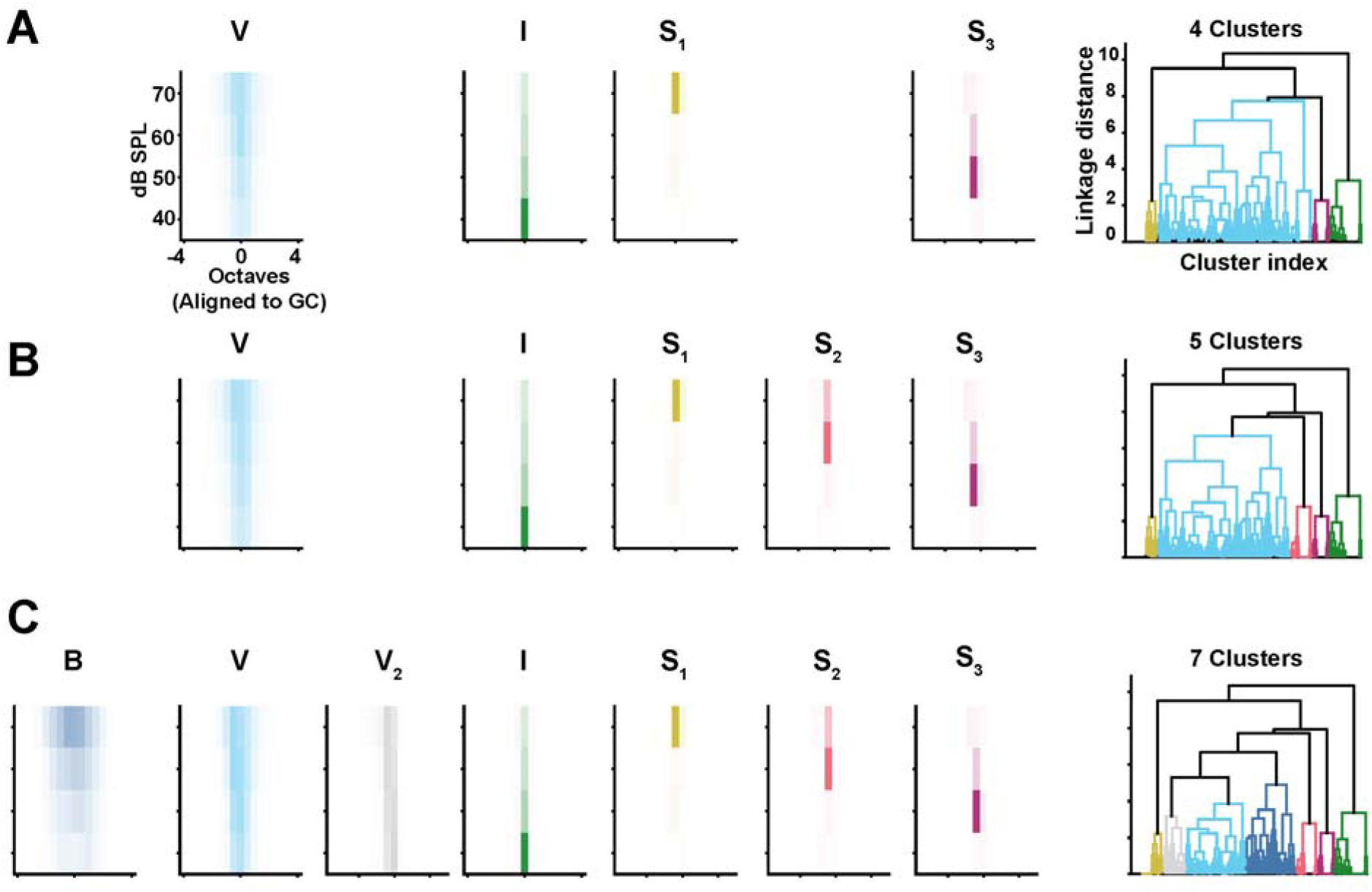
Unsupervised clustering of L6 CTNs with varied number of clusters. A) Left: Averaged FRA maps if only four clusters are considered. Right: Dendrogram with different colors representing different FRA shapes. B-type neurons in (A) and (B) were merged into V-type neurons. S3-type neurons in (A) and (B) were merged into I-type neurons. B) Left: Averaged FRA maps if five clusters are considered. Right: Dendrogram with different colors representing different FRA shapes. B-type neurons in (A) and (B) were merged into V-type neurons. S3-type neurons in (A) and (B) were merged into I-type neurons. C) Left: Averaged FRA maps if seven clusters are considered. Right: Dendrogram with different colors representing different FRA shapes. V-type neurons in (A) and (B) were split into V2-type neurons. Ultimately, six clusters were chosen as it displays distinct FRA shapes without repetition.

**Figure S2:**
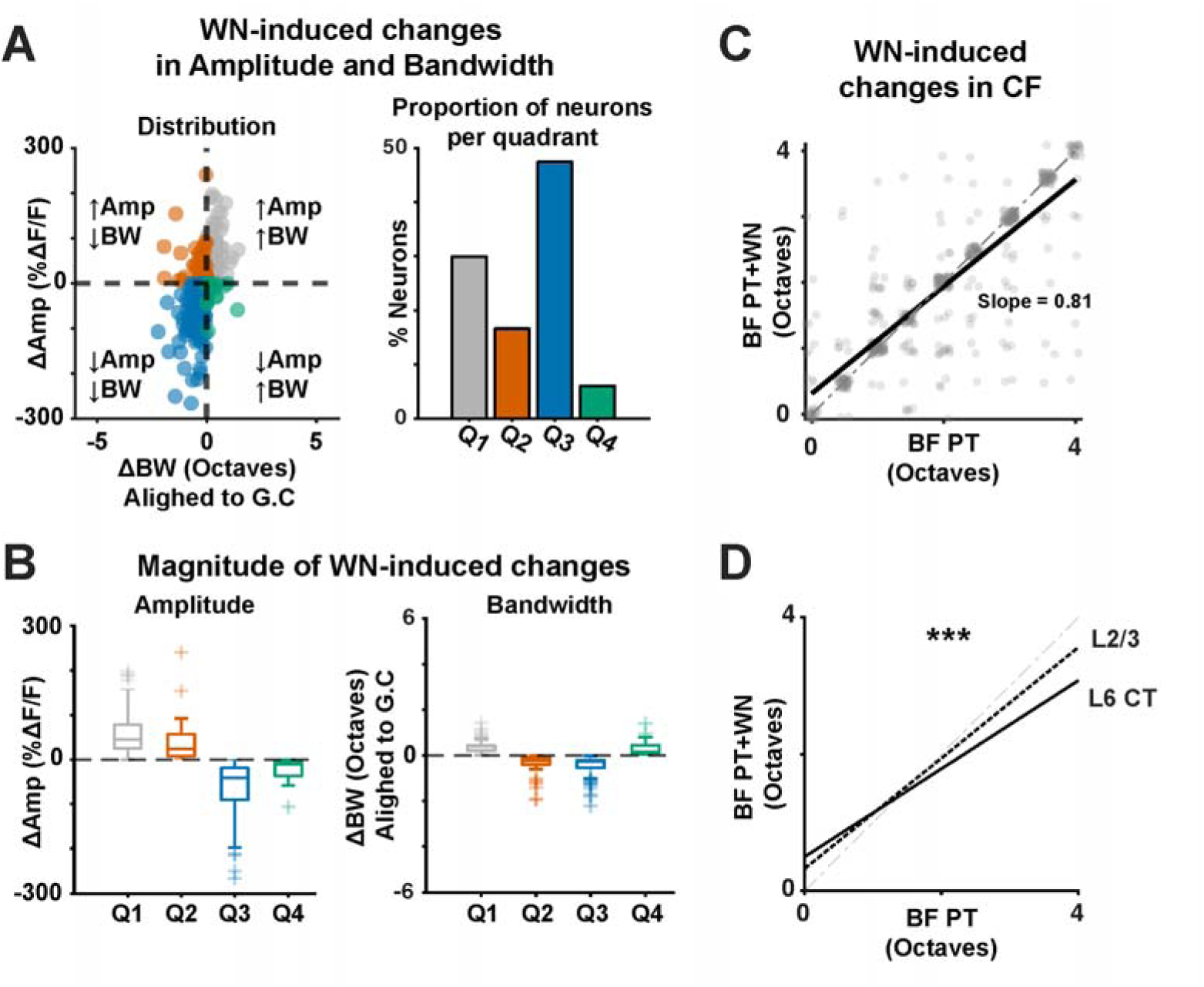
L2/3 neurons show less changes to white noise stimuli than L6 CTNs. (A) Left: Scatter plot of differences in response amplitude (Amp) and bandwidth (BW) as a result of WN stimuli (11 FOVs, 10 mice, 560 sound evoked neurons). (B) Right: Proportion of neurons in each quadrant of the scatter plot. Magnitude of changes in Amp (left) and BW (right) of L2/3 neurons as a result of WN stimuli. (C) Linear regression model between CFs at different experimental conditions (PT and PT+WN). (D) Linear regression model including an interaction term between L2/3 and L6 CTNs and experimental condition (p-value = 8.21 × 10^−5^)

